# Novel Body-Selective Regions Responsive to Bodies Away from the Center of Gaze

**DOI:** 10.1101/2025.06.05.654364

**Authors:** Yu Zhao, Matthew W. Shinkle, Mark D. Lescroart

## Abstract

We report the existence of two previously undescribed body-selective visual regions in the human brain, which we term the Ventromedial Body Area (VMBA) and the Medial Body Area (MBA). We localize these regions based on high-signal localizer contrasts and characterize selectivity in them with encoding model analysis of BOLD fMRI responses to thousands of naturalistic images. These regions respond to images of bodies away from the center of gaze.

## Main Text

Recent advances in neuroimaging, including large high-field datasets^1^ and naturalistic stimuli coupled with sensitive computational analyses, have revealed previously undescribed regions selective for food^2–4^, scenes^5^, words^6^, and numerosity^7^. These new discoveries are critical, because theories of cortical organization^8–11^ rely on accurate and complete estimates of visual selectivity. In contrast to these recent findings in other domains, no new body-selective areas have been described for 14 years^12–15^. To better understand the representation of bodies in natural contexts and to inform theories of cortical organization, we here perform a principled exploration to identify and characterize body-selective regions in the human brain.

To this end, we analyzed data from the Natural Scenes Dataset (NSD), a large dataset of 7T BOLD fMRI responses. To search for candidate body-selective areas, we first analyzed the NSD category localizer data. For each subject, we computed a contrast (an unequal variance t-test) between estimated responses to bodies and limbs versus estimated responses to faces, objects, characters, and places^16^. These contrasts revealed known regions, i.e. the Extrastriate Body Area (EBA) and Fusiform Body Area (FBA), in addition to at least two other regions of cortex with apparent selectivity for human bodies. Based on these contrasts we demarcated two regions for further investigation. The first of these, which we term the ‘Ventromedial Body Area’ (VMBA) was localized as a patch of strong positive contrast for bodies (*t* > 3.83, *p* < 0.05 FDR corrected^17^) posterior to the Parahippocampal Place Area (PPA) in the fork of the Collateral Sulcus. The second candidate region, which we term the ‘Medial Body Area’ (MBA), was also found near a known scene-selective region, the Retrosplenial Complex (RSC), in dorsomedial parietal cortex along the Parieto-Occipital Sulcus. Both regions were found in all eight subjects, with MBA predominantly in the right hemisphere (six right only, two bilateral).

Mapping of these regions from individual subjects onto an average cortical surface (FsAverage^18^) revealed anatomical consistency (Fig. 1c).

**Figure 1.**
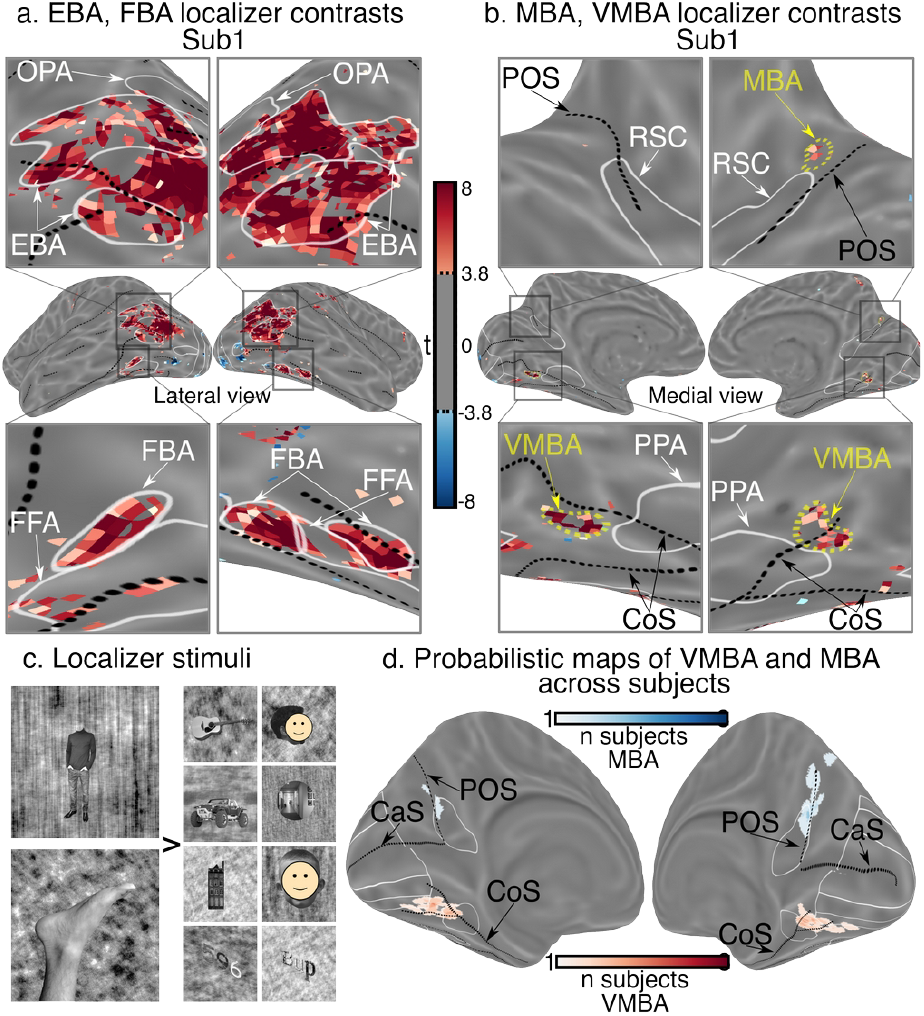
Functional body localizers reveal patches of apparent body selectivity in consistent locations near scene-selective regions. **a**. Body localizer contrast (t test of bodies > all other categories at each voxel) showing known lateral areas **b**. Body localizer contrast showing new ventral and medial areas. See Supplementary Fig. 1 for all subjects. **c**. Sample images shown in localizer experiment **d**. Probability map of locations for MBA and VMBA. These regions fall in consistent locations across subjects, in the fork of the Collateral Sulcus (CoS) and along the Parieto-Occipital Sulcus (POS). MBA is more reliably localized in the right hemisphere.

To investigate whether these regions would respond to diverse bodies in naturalistic contexts, we used voxelwise modeling to analyze the main NSD experiment data. This dataset consists of estimated response to ∼10,000 naturalistic images in each of eight subjects. We extracted a set of body features describing the 2D image locations of bodies, 2D image locations of heads, and the number of bodies in each image (Fig. 2a, see methods for details). We then fit a voxelwise encoding model for each subject. This fitting procedure used the ∼9,000 images unique to each subject as a training set to establish a linear mapping between these body features and fMRI responses in each voxel. We then used the fit model for each subject to predict the responses to the ∼1,000 images shown to all participants in NSD. Predicting independent data with the fit encoding model constitutes a test of the hypothesis that we have accurately captured features that are related to brain responses in each voxel. We found that the body models accurately predicted responses within both VMBA and MBA (Supplementary Fig. 5). To account for naturalistic correlations between body features and other stimulus characteristics, we tested our body model against plausible alternative hypotheses. To this end, we fit additional encoding models based on (1) low-level visual features extracted by spatial Gabor filters and (2) a set of categorical features describing scene-related characteristics. (See online methods for details.) We fit encoding models for each feature set separately, as well as for each combination of feature sets. We applied variance partitioning^19^ to these models to reveal the amount of variance in voxel responses explained uniquely by each model.

**Figure 2.**
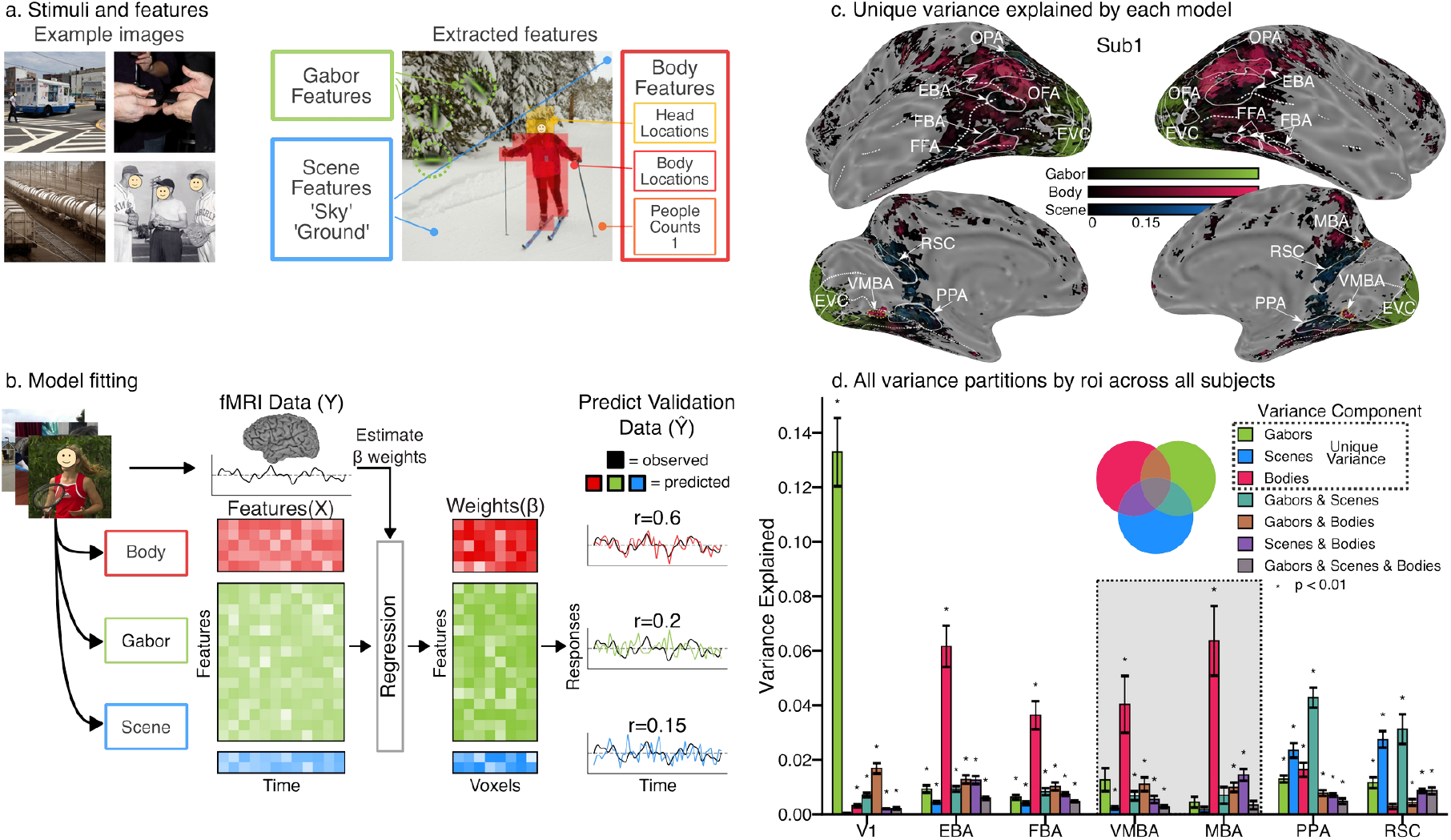
Encoding model procedures. **a**. Stimulus features extracted as the basis for encoding models. **b**. Flow chart of model fitting procedures. Stimulus features and BOLD responses in training data were used to fit weights that reflect the importance of each feature in each voxel in each subject. Fit models were used to predict responses to 1,000 independent stimuli in the validation data set. **c**. Unique variance predicted by each model in one subject. Note that blending of colors, which would indicate multiple models contributing unique variance to a given region, was possible in this plot but not observed. See Supplementary Fig. 2 for all subjects. **d**. Bar plot of unique variance explained by region across subjects. Per c. and d., VMBA and MBA show much more unique variance explained by the body model than by any other set of features, and comparatively little variance is shared between models. Asterisks indicate variance components reliably greater than zero (one-tailed t-tests, p < 0.01, FDR-corrected across all region-component combinations).

We observed significant unique variance explained by our body model within both VMBA and MBA, as well as in established body selective regions EBA and FBA (Fig. 2c, all colored regions *r*^*2*^_unique_ > 0.0088, p < 0.05, FWE corrected^20^). Furthermore, we saw minimal unique variance explained by body features in PPA and RSC, despite their anatomical proximity to VMBA and MBA. In contrast, our scene category model explained substantial unique variance within PPA and RSC, but very little in MBA and VMBA. Our Gabor model explained substantial unique variance in early visual cortex, but little within VMBA and MBA. Taken together, these results argue that MBA and VMBA are functionally distinct from neighboring

ROIs and respond selectively to bodies. Furthermore, these results confirm that responses to bodies in VMBA and MBA cannot be explained by correlations of bodies with low-level image features or structural elements of scenes.

The co-occurrence of bodies and faces in naturalistic stimuli raises the question of whether MBA and VMBA may be selective for faces. To address this question, we averaged regression weights corresponding to different visual field locations for heads and bodies to obtain two values per voxel: the average weight on bodies and the average weight on heads. Fig. 3c shows these values averaged across ROIs and subjects.

**Figure 3.**
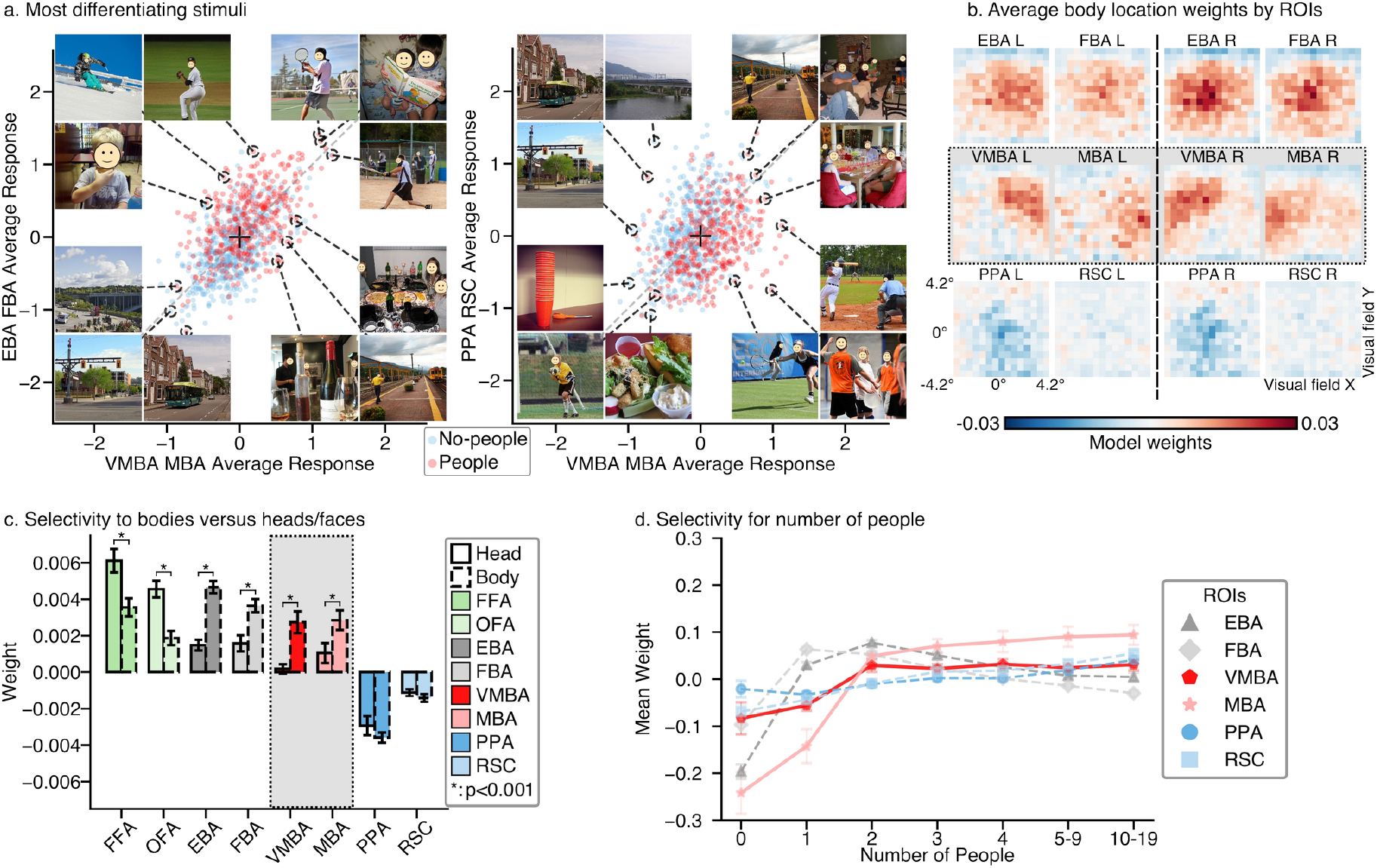
VMBA and MBA selectivity analysis. **a**. Plot of cross-subject average z-scored responses to each of 907 shared images in VMBA and MBA on the x axis versus average response in EBA and FBA (left) or PPA and RSC (right) on the y axis. Red dots correspond to images with people. **b**. Average weights for each of 15×15 body locations for six ROIs in each hemisphere. VMBA and MBA appear selective for bodies away from fixation, while EBA and FBA are selective for bodies near fixation, and scene-selective PPA and RSC have low weights for body features anywhere in the visual field. **c**. Average responses to bodies versus heads by ROI. Known face-selective regions (FFA and OFA) respond more to heads, while known body regions (EBA and FBA)—and VMBA and MBA—respond more to bodies. **d**. Feature weight analysis shows selectivity for number of people in each ROI. VMBA and MBA seem to prefer more people in a scene than do EBA and FBA. See Supplementary Fig. 6 for statistical comparisons between body count bins.

As expected, face-selective ROIs showed higher weights for heads, and conventional body-selective ROIs showed higher weights for bodies (Fig. 3c). Crucially, both VMBA and MBA exhibited significantly greater weights for bodies than for heads (t > 5.4, p < 0.001), with contrast magnitudes comparable to those observed in established body-selective regions EBA and FBA (t > 6.3, p < 0.001). Thus, these areas are approximately as body-selective as EBA and FBA.

We then asked whether VMBA and MBA differ in their body feature selectivity compared to other body-selective regions. First we visualized images that elicited high or low responses in each region compared to other regions. Fig. 3a shows that VMBA and MBA consistently responded to images with people, as did EBA and FBA. However, the new regions preferred multiple people slightly away from fixation, whereas EBA and FBA preferred single people near the fovea. We also quantified selectivity of these regions for both body location and the number of bodies present in each scene. We first examined body location weights from our joint encoding model fit. The weights of this model describe the relationship between the responses of a voxel and the presence or absence of bodies in different image locations.

Thus, we visualized the average weights for each region to examine the pattern of body location selectivity over image space for each region. Consistent with the qualitative picture from Fig. 3a, encoding model weights for VMBA and MBA (Fig. 3b) revealed a preference in both regions for bodies located away from the fovea, while EBA and FBA displayed clear preference for foveally located bodies. This is consistent with previous work showing selectivity for central (versus left/right) bodies in EBA^21,22^.

To quantify the preference of each area for number of people, we examined encoding model weights for our ‘body counts’ feature set. Fig. 3d shows the average weight assigned to each number of people in VMBA, MBA, and other reference regions. Both VMBA and MBA responses increase more sharply between one and two people than do responses in EBA and MBA. PPA and RSC, in contrast, showed little change in responses across number of people present.

Taken together, these analyses of selectivity revealed a clear functional distinction of VMBA and MBA from known body-selective (EBA and FBA) and scene-selective (PPA and RSC) regions. VMBA and MBA preferred stimuli containing multiple bodies located away from the fovea. This suggests that VMBA and MBA play a complementary role in visual processing of bodies compared to EBA and FBA. Importantly, VMBA and MBA do not overlap with the reported locations of the recently described ‘scene memory network’^23^. Scene memory regions have been described anterior to PPA and RSC (and OPA), but the new regions reported here are *posterior* to PPA and superior to RSC.

The existence of these regions expands our understanding of body processing beyond the common focus on bodies as foveal stimuli (due to fixation of a face or body). Indeed, there are many natural circumstances in which we do not fixate bodies, e.g. in social situations involving more than one person, in team sports, or while walking through crowds. Under all these circumstances, bodies are assessed not necessarily for their identity or emotions, but for their peri-personal locations, actions, and/or spatial context. Thus, these areas may represent information about the social or spatial context of bodies, or about bodies *as* context for other stimuli (including other bodies). This could explain the proximity of these areas to scene-selective areas PPA and RSC. It could also explain the location of MBA near the precuneus, which has been implicated in visual imagery, episodic memory, and direction of attention among many other functions^24^.

These regions may have been overlooked because they are small and may not be precisely aligned in common cross-subject template spaces, because category localizer stimuli commonly only include one body at a time, and/or because many studies collect only a few localizer runs and thus do not have as robust a signal as do the localizers in NSD. We suggest that there may be many such overlooked regions in the brain that are small and narrowly selective.

## Online Methods

## Feature Sets

### Body Head Locations Features

To extract information about the locations of bodies within the NSD stimuli, we combined two types of annotations from the source COCO dataset: instance segmentation masks and body keypoints. We used the non-crowd instance segmentation masks for the ‘person’ supercategory. These masks delineate the pixels occupied by human bodies, while the body keypoint annotations provide locations of anatomical landmarks on each person, such as eyes, nose, shoulders, and knees.

We used these complementary annotations to partition the person masks into two feature sets, one for bodies and another for heads. This partitioning served two purposes. First, it made the body location features consistent with the body stimuli used in the category localizers (such that neither contained faces or heads). Second, it enabled subsequent analyses to distinguish between selectivity for bodies versus heads or faces.

To achieve this partitioning, we leveraged the body keypoint annotations. We found the nearest keypoint to each pixel within ‘person’ masks for all stimulus images. Each pixels was labeled as ‘body’ if the nearest keypoint was a body keypoint (e.g., shoulder, elbow, knee), and as ‘head’ if the nearest keypoint was a head keypoint (e.g., nose, eye). While body keypoints are present for most person instance masks, there are some masks for which these annotations are absent. Consequently, some very small or distant bodies are not captured in this feature set.

Masks were defined at the original resolution of the COCO images and cropped in the same fashion as the NSD stimuli. Finally, to reduce the dimensionality of the body and head feature maps, we spatially downsampled them to a size of fifteen by fifteen using inter-area interpolation. This process resulted in two fifteen-by-fifteen feature spaces, each containing continuous values ranging from zero to one, indicating the presence of bodies or heads at each location. We applied a threshold of 0.2 to binarize these continuous values, such that all values in the body feature space were zeros and ones.

### Gabor Features

To quantify low level visual features that may correlate with body characteristics, we generated a set of Gabor features for each image, using a method identical to the static Gabor feature space described in Nishimoto et al., 2011^25^and implemented in pymoten (https://github.com/gallantlab/pymoten) This entailed first converting each cropped image to luminance values and the multiplying each image by Gabor filters varying in size, position, orientation (0, 45, 90 and 135 degrees) and spatial frequency (0, 2, 4, 8, 16 and 32 cycles/image). This resulted in 1,425 features describing patterns of luminance and orientation variation in the NSD stimuli.

### Body Counts Features

We sought to model selectivity to the number of human bodies present in an image, and to control for this factor in examination of our other models. Therefore, we constructed a ‘body counts’ feature set, describing the number of human bodies in each image. First, we used the same ‘person’ instance masks from the COCO dataset used in creating the body and head locations features. We excluded any masks containing the COCO label ‘is_crowd’ (a tag indicating that a given mask is for a crowd, which contained an unspecified number of bodies). We also excluded each mask which lay entirely outside of the portion of the image remaining after the NSD cropping. We then computed the count of remaining masks for each image.

Regression models fit directly on these counts would only be able to capture linear relationships between the number of bodies and BOLD responses. In order to allow estimation and interpretation of potential non-linear selectivity to body counts, we converted this single continuous feature into six binned (binary) features. Specifically, we binned the number of bodies in each image using the following bins: 0 bodies, 1 body, 2 bodies, 3 bodies, 4 bodies, 5 to 9 bodies, and 10 to 18 bodies. Because these bins are non-overlapping and include the full range of body counts in the NSD stimuli, this results in a one-hot vector for each image.

### Scene Categories Features

In naturalistic stimuli, bodies and their locations may correlate with various features of scenes. To distinguish between selectivity to bodies and spurious selectivity due to correlations with scenes, we defined an additional set of binary ‘scene categories’ features based on the following COCO ‘stuff supercategories’: ‘floor’, ‘structural’, ‘wall’, ‘ceiling’, ‘window’, ‘building’, ‘water’, ‘sky’, and ‘ground’. For each supercategory in each image, we included only scene labels for which the image mask for the supercategory label fell partially within the NSD crop box for the image. This resulted in 9 binary features indicating the presence or absence of each scene feature.

### BOLD Data

We analyzed two sets of BOLD data from the Natural Scenes Dataset (NSD)^1^: category localizer contrasts and estimated betas from the main NSD experiment. Both datasets were acquired using 7T fMRI from eight subjects viewing images. We describe these briefly below, but for full details of the original data collection and preprocessing, see ^1^.

### Functional Localizers and Region of Interest Definition

Regions of interest were defined based on separate functional localizer experiments conducted on NSD subjects. Category-selective regions were localized following the protocol in ^16^. Briefly, subjects viewed grayscale images from different stimulus categories in 4-second trials. Per recommended use of these localizer stimuli (http://vpnl.stanford.edu/fLoc/^16^) we used an unequal-variance *t* contrast of bodies (including headless bodies and limbs) versus all other categories (objects, i.e. cars and instruments; places, i.e. houses and corridors; faces, i.e. adult and child faces; and characters, i.e. words and numbers) to identify body-selective regions, including the Extrastriate Body Area (EBA), the Fusiform Body Area (FBA), and our two new regions, the Ventromedial Body Area (VMBA) and the Medial Body Area (MBA).We used a contrast of places versus all other categories to define scene-selective areas including the Parahippocampal Place Area (PPA), the Occipital Place Area (OPA), and the Retrosplenial Complex (RSC, also called the Medial Body Area or MBA). We used a contrast of faces versus all other categories to define face-selective areas including the Fusiform Face Area (FFA) and the Occipital Face Area (OFA). There were two non-contiguous sub-sections of FFA apparent in most subjects, but since our purpose was not to study subtleties of face recognition *per se*, these regions were combined into one region.

For all category-selective regions, we hand drew regions of interest boundaries based on un-thresholded functional contrasts using pycortex (http://github.com/gallantlab/pycortex^26^). We drew boundaries at locations of rapid change in the relevant contrast value. In practice, this selected regions of reliable statistical significance per the thresholded contrast maps (see Fig. 1). In rare cases, contrasts for two categories were significant over the same set of voxels.

For example, a region at the border of FBA and FFA for some subjects was reliably activated for face versus other and also for body versus other. In such cases, we drew boundaries that divided any overlap evenly between the two adjacent regions such that the regions did not share any common voxels. This is a pragmatic decision rather than a strongly theoretically based decision. We argue that it is more principled than the alternatives, i.e. having data from the same voxels contribute to multiple bars in the same bar plot, or excluding voxels that *are* functionally selective from analysis.

We defined V1-V3, hV4, V3A, V3B, IPS0, LO1, and LO2 based on retinotopic population receptive field (pRF) mapping that was also conducted for NSD subjects. Boundaries for these regions were the same as regions provided with the NSD dataset. Since it is well established that motion-selective regions exist anterior and adjacent to LO2, we defined an additional region in this area based on multiple criteria. The most common way to define motion-selective regions MT and MST are with localizers for expanding versus stationary dots^27^. No such localizers were collected for NSD participants. Thus, we relied on other criteria that have been ascribed to these areas. Specifically, MT and MST are associated with retinotopic regions TO1 and TO2^28^, are known to fall in a posterior continuation of the Inferior Temporal Sulcus (ITS)^29^, and are surrounded by body-selective regions on at least 3 sides^22^. Thus, we drew a single region (hMT+, a common term for the undifferentiated areas MT and MST) that was adjacent and anterior to LO2, that spanned two visual field maps in the extant pRF data, that fell within a posterior continuation of the ITS, and that showed minimal activation in the body versus other category contrast. We acknowledge this is not a field-standard way of defining hMT+, but we argue that an attempt to define hMT+ based on alternative criteria is more in keeping with past literature than ignoring the existence of a well-established region in this area.

We provide two further notes on our ROI definitions. First, we observed another body-selective region near the posterior end of the Intraparietal Sulcus and partly overlapping IPS0 in some subjects. This region has been reported in several previous papers. We did not include this region in the present analyses, since (a) it is clearly anatomically distinct from the present regions of interest (VMBA and MBA), (b) its function is less well established than that of EBA and FBA, (c) it was less reliably identifiable across participants in the localizers than were VMBA and MBA, and (d) it was not well predicted by our body model. Second, we note that one of the regions we describe (VMBA) partially overlaps with the peripherally selective part of retinotopic visual areas VO1 and VO2 in the Wang atlas^30^ (i.e. in average-brain cortical space). These areas have not been previously reported to be body selective, and are overall biased toward foveal representations^31^, and we could not confidently identify VO1 and VO2 in individual brains. Thus, this overlap may be an artifact of imperfect cross-subject alignment.

However, since NSD stimuli were of necessity presented at a relatively small visual angle (8.4^°^x8.4^°^), further studies will be needed to assess overlap of the representation of bodies in our newly described areas with VO1 and VO2 in the visual periphery.

### Naturalistic Images

In the main NSD experiment, subjects viewed color natural scenes while performing a continuous recognition task. Each subject was presented with 8,302-9,000 distinct images two to three times each across 30-40 scan sessions. A separate set of 1,000 images was shown to all subjects, though not all subjects saw all images (four subjects saw 1,000 images, two saw 930, and two saw 907).

Our analyses used the version 3 (b3) beta estimates at the native 1.8 mm isotropic resolution in individual subject space. To ensure data quality, we included only voxels meeting a noise ceiling threshold of *r*^*2*^=0.2. We divided this set of data into a training set (unique images for each subject) and a test set (∼1,000 shared images). This division allowed us to train models on a large, diverse stimulus set while reserving a common set for cross-subject validation. All modeling data (i.e. the NSD main experiment) was independent of the data used to define the regions (i.e. the NSD localizers).

### Regression Fitting

We used regularized linear regression to estimate predictive relationships between features from one or more feature sets and BOLD responses in each voxel. For feature preprocessing, we applied z-scoring only to the Gabor features to account for different possible ranges of values across different Gabors. All other feature sets (Body Locations, Head Locations, Body Counts, and Scene Categories) were left in their original scale (0-1). Estimated responses for the NSD data were not normalized prior to regression fitting.

We used the Himalaya Python package^32^ for model fitting. To determine the optimal regularization parameters for each voxel and feature set, we used cross-validation with a scaled L2 penalty on model weights. All models were fit with an intercept term.

In addition to fitting models for individual feature sets, we also performed joint fits incorporating multiple feature sets using banded ridge regression. This approach serves two purposes. First, it enables variance partitioning, quantifying the unique explanatory contribution of each set of features to predictions of a voxel’s response. Second, by fitting alongside other feature sets with which body features are likely to be correlated, joint fitting reduces the effects of these correlations on the body model weights.

We computed seven total fits for each subject: one for each individual feature set (Gabor, Scene Categories, and a Body composite including Body Locations, Head Locations, and Body Counts), and one for each combination of these sets.

### Statistical Significance Testing

To assess the statistical significance of the variance explained by our encoding models, we performed permutation testing using the COPTeRR (Compute-Optimized Permutation Testing for Ridge Regression) package (https://github.com/piecesofmindlab/copterr). COPTeRR enables computationally efficient permutation testing on ridge regression models by avoiding model refitting for each permutation, instead leveraging the closed-form solution of ridge regression to directly compute model weights and performance metrics from permuted data. For each subject, we generated 10,000 permutation indices using a block permutation scheme with a block length of five time points to preserve temporal structure in the data while breaking the relationship between features and brain responses. Permutation indices were generated separately for training and validation data using independent random seeds to ensure proper cross-validation. For each permutation, we computed correlation coefficients between predicted and actual brain responses using the permuted feature-response mappings. Statistical significance was determined by comparing the true model performance to the null distribution of permuted performance values, with significance thresholds calculated at α = 0.05 using the 95th percentile of the permutation distribution. To control for multiple comparisons across voxels, we applied family-wise error rate (FWE) correction by taking the maximum correlation coefficient across all voxels for each permutation and using the 95th percentile of these maxima as the significance threshold^20^.

