## Supplementary Figures for "Novel Body-Selective Regions Responsive to Bodies Away from the Center of Gaze"

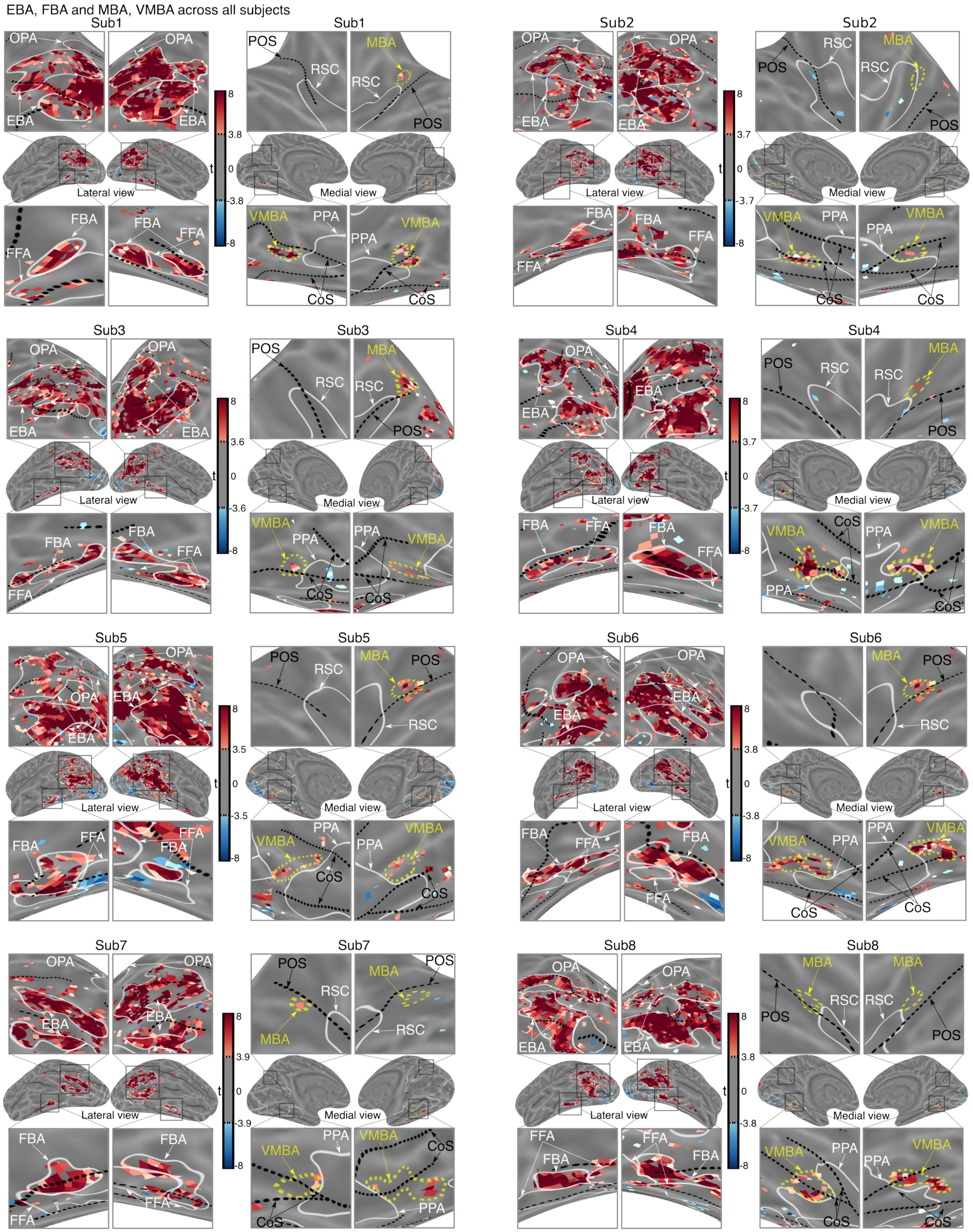


*Supplementary Figure 1: Body localizer contrast (bodies > objects) for all subjects, showing both known lateral body-selective regions and the newly identified ventral and medial body areas. Each subject's t-map is thresholded at $p < 0.05$, FDR corrected, for visualization.*

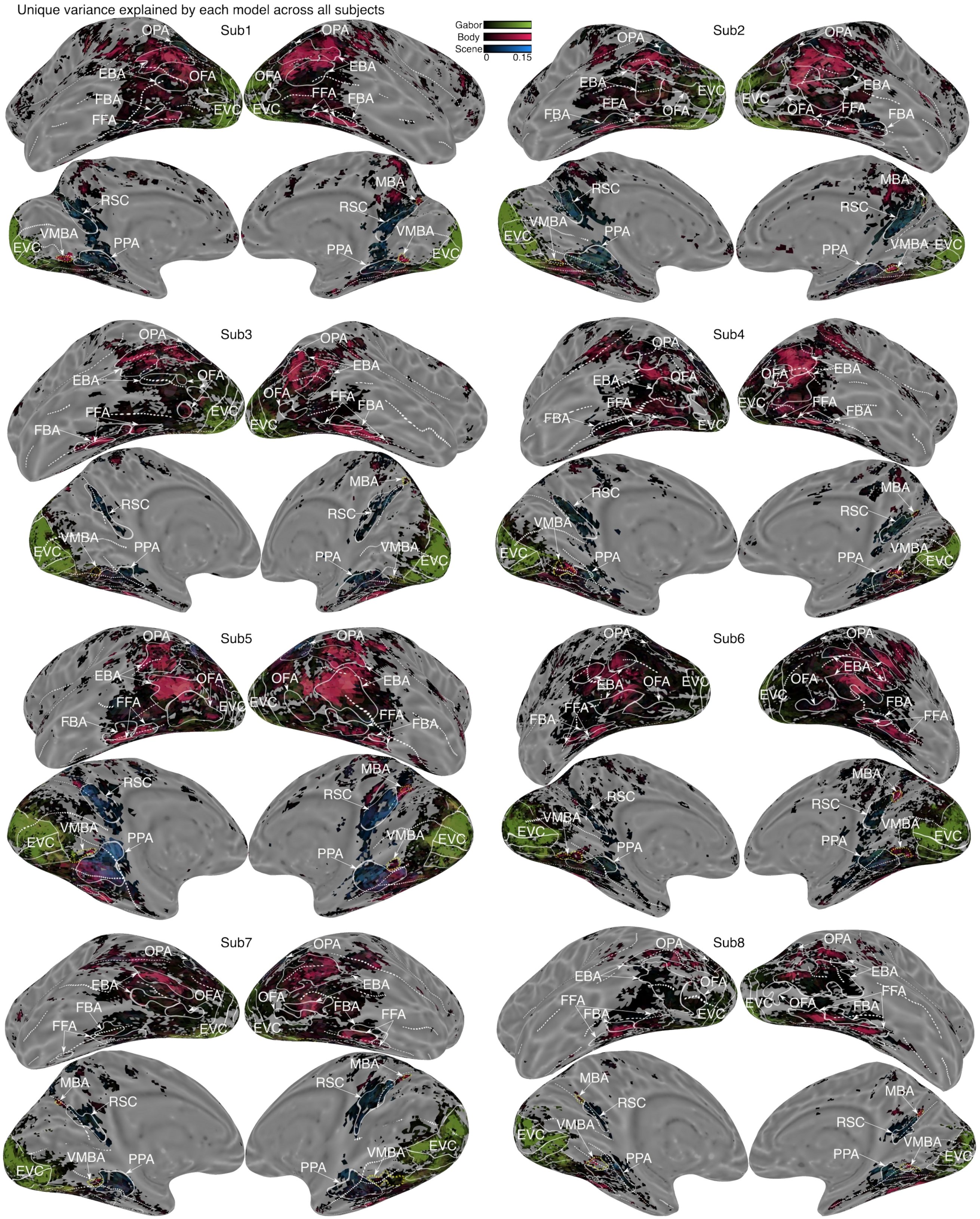


*Supplementary Figure 2: Unique variance explained by each encoding model across all subjects. VMBA and MBA consistently showed strong unique variance explained by the Body model across subjects.*

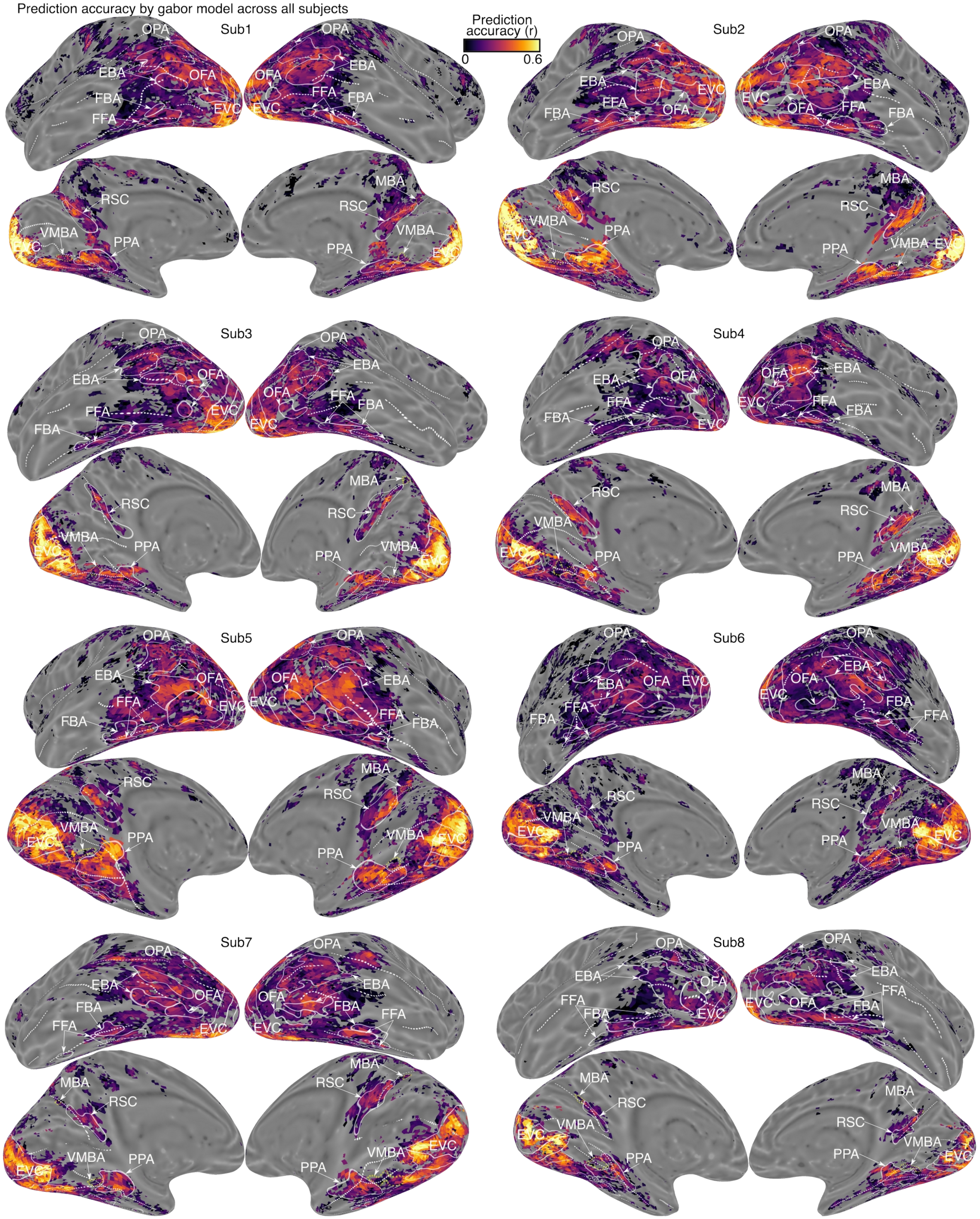


*Supplementary Figure 3: Prediction accuracy (r values) of the Gabor model across all subjects. Maps show the voxelwise Pearson correlation between predicted and observed BOLD responses in the independent validation set. See Fig. 2b for model fitting schematic.*

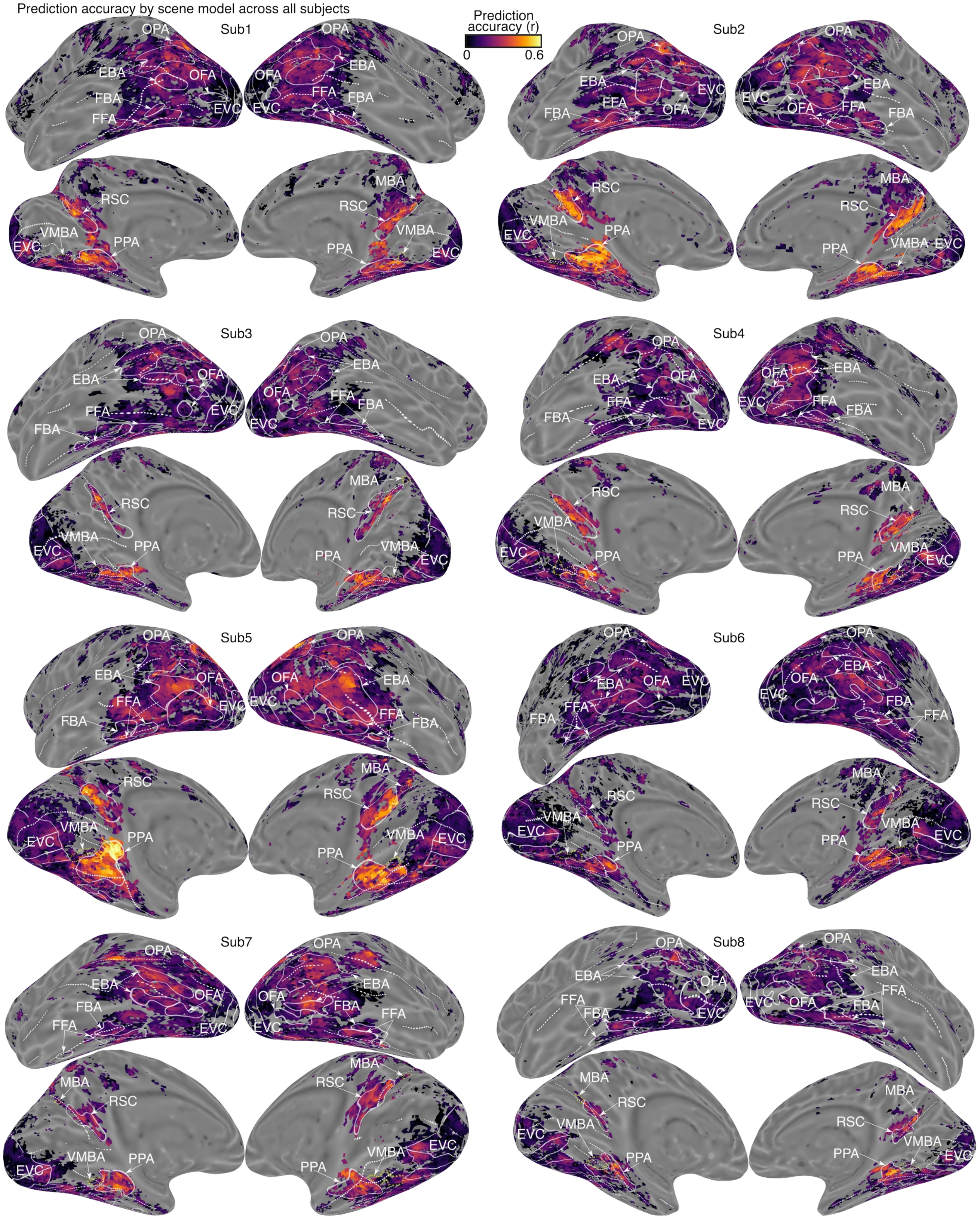


*Supplementary Figure 4: Prediction accuracy (r values) of the Scene model across all subjects.*

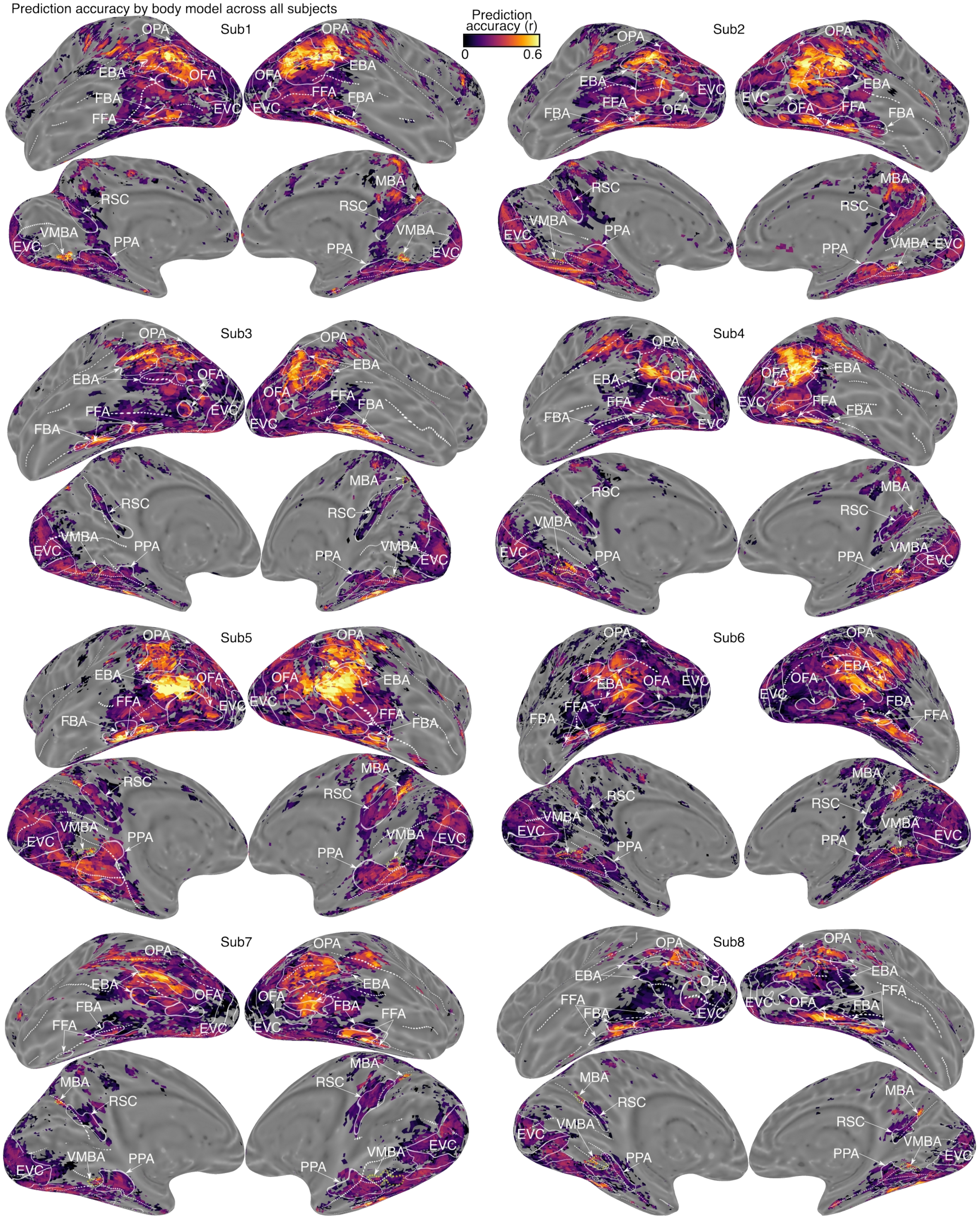


*Supplementary Figure 5: Prediction accuracy (r values) of the Body model across all subjects.*

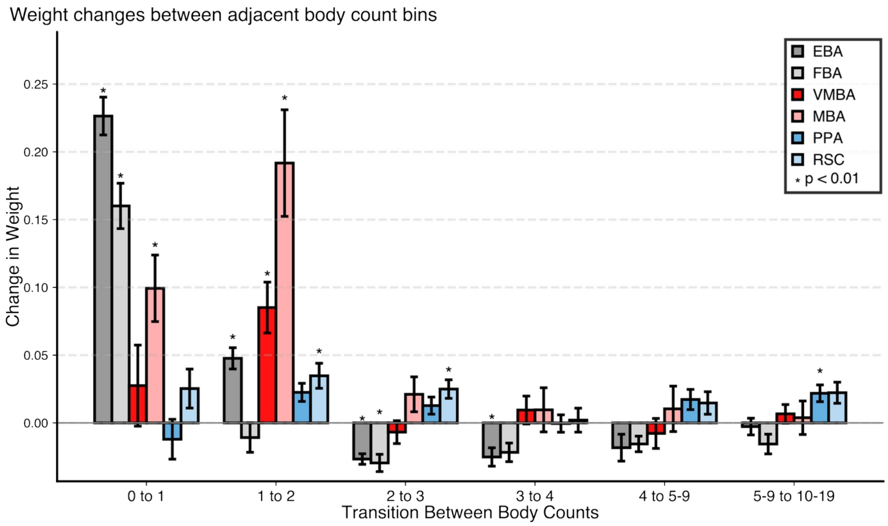


*Supplementary Figure 6: Body count transition analysis reveals differential response patterns across brain regions to changes in person number. EBA, FBA exhibit the strongest weight increases from zero to one person. Conversely, VMBA, MBA show peak increases from one to two people, highlighting sensitivity to single-to-multiple person transitions. Error bars represent SEM across subjects; asterisks indicate significance from one-sample t-tests against zero ($p < 0.01$, FDR-corrected).*
